# Does it pay to pay? A comparison of the benefits of open-access publishing across various sub-fields in Biology

**DOI:** 10.1101/2022.12.10.519925

**Authors:** Amanda D. Clark, Tanner C. Myers, Todd D. Steury, Ali Krzton, Julio A. Yanes, Angela Barber, Jacqueline L. Barry, Subarna Barua, Katherine M. Eaton, Devadatta Gosavi, Rebecca L. Nance, Zahida H. Pervaiz, Chidozie G. Ugochukwu, Patricia Hartman, Laurie S. Stevison

## Abstract

Authors are often faced with the decision of whether to maximize impact or minimize costs when publishing the results of their research. For example, to potentially improve impact via increased accessibility, many subscription-based journals now offer the option of paying a fee to publish open access (i.e., hybrid journals), but this solution excludes authors who lack the capacity to pay to make their research accessible. Here, we tested if paying to publish open access in a subscription-based journal benefited authors by conferring more citations relative to closed access articles. We identified 146,415 articles published in 152 hybrid journals in the field of biology from 2013-2018 to compare the number of citations between various types of open access and closed access articles. In a simple generalized linear model analysis of our full dataset, we found that publishing open access in hybrid journals that offer the option confers an average citation advantage to authors of 17.8 citations compared to closed access articles in similar journals. After taking into account the number of authors, journal impact, year of publication, and subject area, we still found that open access generated significantly more citations than closed access (*p* < 0.0001). However, results were complex, with exact differences in citation rates among access types impacted by these other variables. This citation advantage based on access type was even similar when comparing open and closed access articles published in the same issue of a journal (*p* < 0.0001). However, by examining articles where the authors paid an article processing charge, we found that cost itself was not predictive of citation rates (*p* = 0.14). Based on our findings of access type and other model parameters, we suggest that, in most cases, paying for access does confer a citation advantage. For authors with limited budgets, we recommend pursuing open access alternatives that do not require paying a fee as they still yielded more citations than closed access. For authors who are considering where to submit their next article, we offer additional suggestions on how to balance exposure via citations with publishing costs.

## Introduction

Ensuring global access to published research is a major goal of scientists today. To achieve this goal, journals have begun to shift toward open access (OA) publishing and away from subscription-based, or closed access, publishing. As OA publishing has grown, several OA publishing modalities have emerged, each offering distinct benefits to authors and readers. The “gold” OA category describes articles made freely available upon publication directly from the publisher’s website under an open license. While gold OA makes accessing research easiest for readers, to publish under the gold OA category, authors must pay an article processing charge (APC). APCs typically place a significant financial responsibility on the author, and depending on available funding, may restrict the number of outlets where they can publish their research.

Gold OA articles may be published either in journals that are entirely open access or in “hybrid” journals, which are traditional, subscription-based outlets that have an option for authors to make articles freely available via payment of an APC (i.e., “hybrid gold”). Hybrid models have the appeal of allowing authors to publish in well-known high impact journals, while simultaneously making them open to non-subscribers. With the rise in OA mandates by funding agencies and universities, the number of subscription-based journals that have introduced a gold OA option has exploded over the past 15 years [Jahn et al., 2022, Björk, 2017]. This increase in hybrid journals has led to an increase in Other gold OA articles published in hybrid journals from an estimated 8,000 in 2009 to 45,000 in 2016 [Björk, 2017]. Furthermore, Elsevier reported a doubling in the number of hybrid gold OA articles published in their hybrid journals every year between 2015 and 2019 [Jahn et al., 2022]. During this same period of time, APCs have increased dramatically, generally outpacing the rate of increases in journal impact and the rate expected if APCs were indexed to inflation [Khoo, 2019]. For example, in a sample of biology journals, 2022 APCs ranged from $1,395 to $5,790 (Table 1).

**Table 1.** Gold OA journal APC changes over time. A selection of gold OA journals from previous study [Krzton, 2019] are listed with the APC values from that study compared to current APC values at the time of our study, representing changes over a 4 year time period. All amounts are listed in US currency. None of these journals were used in the current study, which targeted hybrid access journals, but instead represent general trends in APC changes over time.

| Journal | 2019 APC | 2022 APC |
| --- | --- | --- |
| Genome Biology | 3490 | 5030 |
| Nature Communications | 5200 | 5790 |
| PLoS Biology | 3000 | 5300 |
| Scientific Reports | 1790 | 2090 |
| Database: The Journal of Biological Databases and Curation | 1680 | 2475 |
| Frontiers in Plant Science | 2950 | 2950 |
| Ecology and Evolution | 1950 | 2200 |
| PeerJ | 1095 | 1395 |

With increased pressure to make research open, and rising APCs, authors are left with difficult decisions when choosing how and where to effectively communicate their science. Due to the recent rise of OA publishing, many fully OA journals tend to be younger and lack the well-established audiences of traditional subscription-based journals that have introduced OA options. The generally higher impact of hybrid journals has allowed them to charge higher APCs than fully open journals [Asai, 2022]. While traditional considerations of journal impact and target audience remain important, authors must factor budget into their decisions more heavily now than they did in pre-OA times.

Other OA categories in addition to gold OA have emerged that do not require fees of the authors. Bronze OA describes articles that are designated OA by the journals themselves and at no cost to the author [Piwowar et al., 2018]. However, the process by which articles are selected for bronze OA is unknown and may not be permanent [Piwowar et al., 2018]. A perhaps under-utilized alternative to publishing articles OA in either fully OA or hybrid journals is “green” OA, in which authors self-archive their work by uploading preprints to servers like bioRxiv or by depositing post-prints in institutional repositories or other archives [Tennant et al., 2016, Gadd and Troll Covey, 2019]. Although green OA is subject to journal permissions, formatting restrictions, and embargo periods, there is no cost to the author under this model, making green OA a particularly appealing alternative to costly APCs.

For certain budgets, the main question authors face when deciding to pay an APC is whether increased access to their work would translate to increased citations. Thus far, attempts to answer whether OA publishing confers a citation advantage to authors relative to publishing closed access have produced mixed results. While many studies have found support for an OA citation advantage, others have found the opposite [Dorta-González et al., 2017]. Furthermore, studies that have found support for a citation advantage between OA and closed access [Piwowar et al., 2018, Sotudeh et al., 2019, Ottaviani, 2016] have been careful to avoid concluding that OA status leads to greater citations due to methodological and statistical challenges involved in designing a robust citation study that limits the impact of confounders, including language [Lewis, 2018, Basson et al., 2021, Moed, 2007]), field or subject area [Archambault et al., 2016, Holmberg et al., 2020, Hubbard, 2017], and journal age. Therefore, any attempt to estimate differences in citation rates between access types must be aware of the potentially confounding forces that may influence citations and account for article attributes that may influence citation rates.

Hybrid journals provide the closest thing to a direct comparison that could be used to test whether a citation advantage for OA publishing exists [Björk, 2017, Harnad and Brody, 2004, Tang et al., 2017]. Specifically, hybrid journals circumvent the confounding factor of variation in journal impact as both OA and closed access articles can be compared for the same journal. However, few existing studies have taken advantage of the comparison presented by hybrid OA journals to test if OA confers a citation advantage. One such example recovered evidence that OA articles were cited earlier and more frequently than closed access articles published in the same journal during the same period of time [Eysenbach, 2006]. However, a comprehensive assessment testing whether OA confers greater citations while taking differeneces among subject areas into account is lacking.

Biology has a higher than average number of hybrid OA papers than other fields [Laakso and Björk, 2016], and a citation advantage for OA has been documented [Archambault et al., 2016, McCabe and Snyder, 2014]. Although a few previous studies have looked at the citation pattern in biology, they have largely been limited to just one sub-field within biology [AlRyalat et al., 2019], analyzed relatively small number of records or subset of publications (~3,500 records; [Tang et al., 2017]), or a combination of these [Calver and Bradley, 2010, Clements, 2017]. Furthermore, the results of these studies are conflicting with regard to whether paying to publish OA actually confers a citation advantage to the authors, with some finding a benefit [Tang et al., 2017, Clements, 2017], and others recovering a minimal effect [Calver and Bradley, 2010]. Clements [2017], controlling for self-citation, impact factor, number of authors, and article type, investigated the citation patterns and found an OA citation advantage in three marine ecology journals, yet no such advantage was identified in six conservation biology journals [Calver and Bradley, 2010]. Due to the limited scope of prior research, it remains unclear whether there is any citations advantage provided by OA across sub-fields in biology.

In this study, we addressed the question of whether authors across sub-fields in the biological sciences can expect to gain more citations by paying an APC to publish OA in a hybrid journal. Using the Web of Science database, we collected a sample of 146,415 articles published in 152 hybrid journals published between 2013 and 2018 to compare the rates of citation between OA and non-OA articles. We used these data to assess (1) the degree to which OA articles published in hybrid journals are cited more than non-OA articles, (2) the contributions of factors such as author count, journal impact factor, and sub-field to citation rates, and (3) if and how these factors influenced any differences in citations rates among access types. Based on our results, we provide specific and concrete recommendations to authors that should aid decision-making regarding when to and whether it is worthwhile to pay an APC to publish OA in hybrid biology journals. Overall, our results show a general citation advantage for OA over closed access, and a clear advantage for hybrid gold OA over other types of OA, but this advantage varies depending on article attributes, such as number of authors or journal impact.

## Materials and Methods

Our methodology to acquire and curate the data is laid out in Figure 1. We used Clarivate Analytics Web of Science to obtain bibliographic data from hybrid journals. We selected journals from 12 Web of Science categories encompassing biology: Biochemistry and Molecular Biology, Cell Biology, Entomology, Evolutionary Biology, Genetics and Heredity, Marine and Freshwater Biology, Microbiology, Mycology, Neurosciences and Neurology, Oncology, Plant Sciences, and Zoology. To select only journals with a hybrid publishing model, we excluded all journals that did not include records classified as “Other gold”, the Clarivate Analytics Web of Science designation for hybrid gold articles defined as articles with Creative Commons licenses that are not published in solely OA journals. We also filtered results to remove records not published between 2013 and 2018 and manually verified whether each journal met the hybrid publishing model requirement (Fig. 1). In addition to the bibliographic data obtained for each article (i.e., number of authors and OA status: closed access, bronze, green, or hybrid gold), we collected data for the following journal-level citation metrics from Clarivate Analytics Journal Citation Reports (JCR): JCR Quartile within a selected Web of Science Category, and Article Influence Score (AIS), which quantifies the average influence of a journal’s article within five years of publication. We also collected APCs as of June 2021 from publisher websites for each journal.

**Figure 1.**
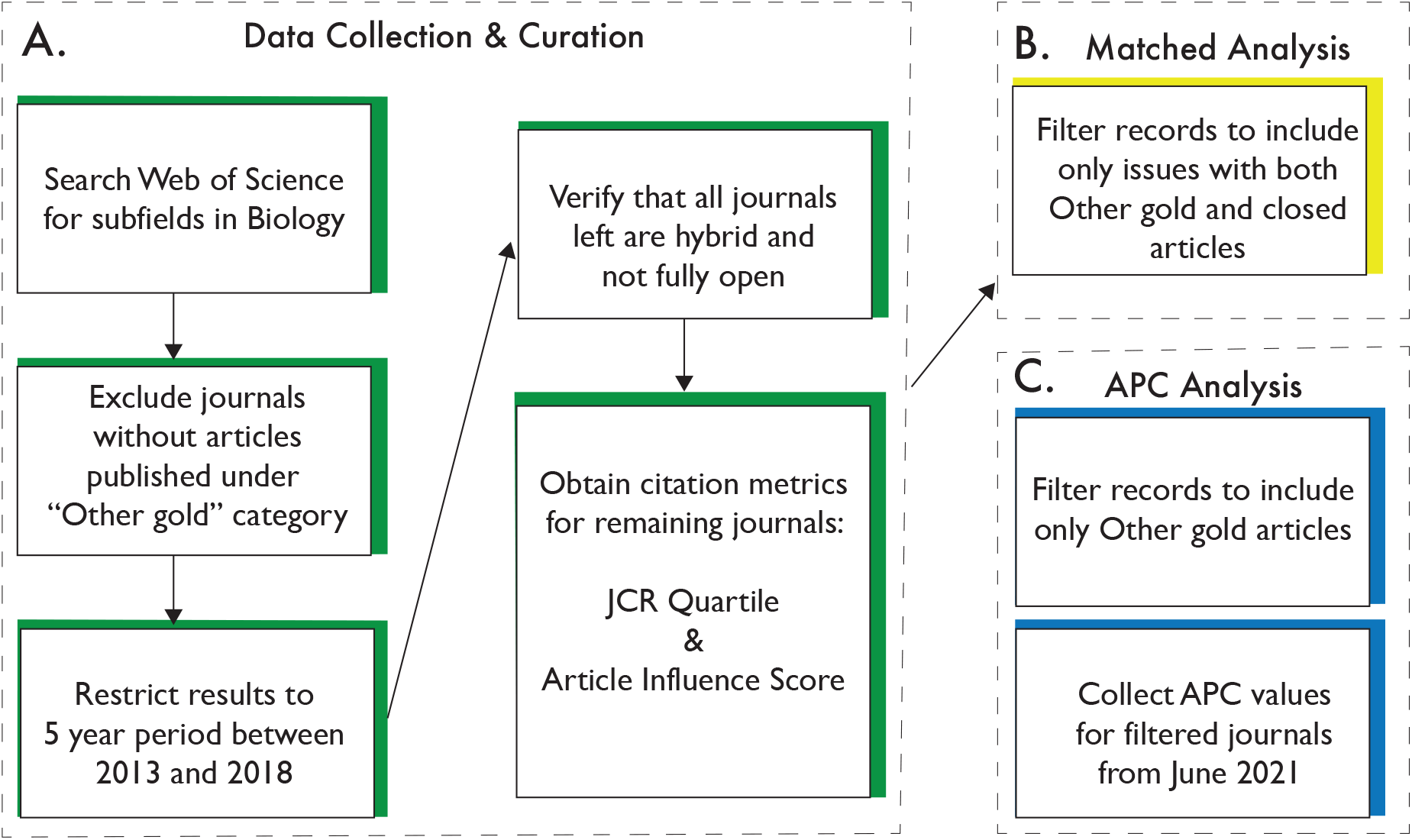
Data preparation. **A.** We obtained citation records for articles published in hybrid journals by conducting searches on Clarivate Analytics Web of Science for different sub-fields in biology. We verified whether the journals found by each search were hybrid journals and, if so, we downloaded data, including number of citations, OA type, etc., for all articles published in each journal between 2013 and 2018. We also obtained citation metrics that we used as predictors in our full and “matched” analyses. **B.** For the matched analysis, we restricted our dataset to compare hybrid gold and closed access articles published in the same volume and issue of our journals. **C.** Lastly, we obtained values as of June 2021 for each journal’s APC to test APC values were associated with number of citations.

To examine the relationship between OA and citation rates while controlling for other factors, we used generalized linear models. In all models, our response variable was raw citations counts. As citation count is likely non-normally distributed, we initially fit generalized linear models to the data with a Poisson distribution for the response. However, likelihood-ratio (Chi-square) tests always indicated that the negative binomial distribution described the data better due to variance inflation in the number of citations (all *χ*^2^ > 3, 437, 870, *p* < 0.0001), and thus we used that distribution for all analyses. In the “full analysis”, we included OA status, author count, JCR quartile (1, 2, 3, or 4), AIS, and year as fixed effects, and field and journal (nested in field) as random effects (Table 3). To improve model convergence and adjust for skewed distributions of the independent variables, we scaled AIS and author count. We also included two-way interaction terms between OA status and each of the other fixed effects. Collinearity among fixed effects was generally low as evidenced by low generalized variance inflation factor scores (all < 1.31). Thus, we considered variance inflation not to be an issue in the full model. In all analyses, statistical significance of fixed effects and interactions was assessed via Type II Wald Chi-square tests using the ’Anova’ function from the R package ‘car’ [Fox et al., 2022]. Pairwise comparisons among groups within a variable were assessed by a Wald Z test with a Bonferroni correction using the ‘emmeans’ R package [Lenth et al., 2022]. All analyses were done in R (version 4.2.1; Team [2020a]) and RStudio (version 2022.07.1+554; Team [2020b]).

**Table 2.** Breakdown of records by sub-field in Biology. A summary of the data used in this study, including sub-field, number of journals targeted and total number of articles used. Additionally, the number of articles in the ”matched analysis” are included as well as a break down by access type of each sub-field.

| Research Area | Number of Journals | Number of Articles | Number of Matched Articles | Bronze | Closed Access | Green | Other Gold |
| --- | --- | --- | --- | --- | --- | --- | --- |
| Biochemistry and Molecular Biology | 1 | 11286 | 8624 | 28 | 9799 | 969 | 490 |
| Cell Biology | 4 | 4575 | 7 | 3868 | 10 | 50 | 647 |
| Entomology | 8 | 5777 | 1567 | 380 | 4669 | 445 | 283 |
| Evolutionary Biology | 16 | 16061 | 3007 | 8133 | 4806 | 1186 | 1936 |
| Genetics and Heredity | 6 | 7190 | 336 | 2512 | 1016 | 232 | 3430 |
| Marine and Freshwater Biology | 6 | 8708 | 2805 | 77 | 7812 | 549 | 270 |
| Microbiology | 8 | 21921 | 596 | 18700 | 720 | 581 | 1920 |
| Mycology | 16 | 7099 | 1627 | 1059 | 5116 | 346 | 578 |
| Neurosciences and Neurology | 5 | 6465 | 1366 | 2707 | 1510 | 927 | 1321 |
| Oncology | 5 | 16152 | 1724 | 9250 | 1413 | 898 | 4591 |
| Plant Sciences | 7 | 9928 | 990 | 7005 | 1603 | 144 | 1176 |
| Zoology | 70 | 31253 | 5432 | 4286 | 22643 | 2934 | 1390 |
| Totals | 152 | 146415 | 28081 | 58005 | 61117 | 9261 | 18032 |

**Table 3.** Model parameters for full open access dataset. The various variables included as predictor variables are listed along with random variables in the statistical model of the full dataset. In the “Matched Analysis”, an additional random variable of “volume and issue” was included and the records were subset to only include volumes/issues with both OA and closed access articles. Finally, in the third “APC analysis” model, the predictor variable of APC was added and articles were subset to only include hybrid gold where authors paid an APC, thus removing the predictor of access type. All models shared the same response variable of Citation Counts.

| Predictor Variables | Random Variables | Response Variable |
| --- | --- | --- |
| Access Type | Field | Citation Counts |
| Author Count | Journal in Field |  |
| Journal Citation Reports (JCR) Quartile |  |  |
| Article Influence Score |  |  |
| Year |  |  |

In a separate analysis, we used a paired design to compare the number of citations between articles published hybrid gold OA and closed access within the same volume and issue of a journal. Thus, we filtered the downloaded records including only issues with both hybrid gold and closed access articles (Fig. 1). These matched data were used in a second statistical analysis (hereafter “matched analysis”) with the response variable of citation count; all independent model parameters for this analysis were the same as those used for the full dataset. However, volume (nested within journal nested within field) and issue (nested within volume nested within journal nested within field) were also included as random effects in the matched analysis. Again, we used a generalized linear model with negative binomial family structure to analyze the data.

In a final sub-analysis, we examined the relationship between citation count and APCs (hereafter “APC analysis”). The full data was restricted to only include articles published under the hybrid gold access model (i.e., those in which APCs had been paid; Fig. 1). A generalized linear model with negative binomial family structure was used to model the relationships between citation count (the response variable) and the same independent variables that were used in the full model analysis with the exception that access type was removed (since all articles were published hybrid gold), and APC charge was included.

## Results

After filtering, we obtained citation data for 146,415 journal articles from 152 hybrid journals across 12 fields within biology (Table 2). The number of records per journal averaged 963 and ranged from 15 to 11,286. Records across research fields averaged 12,201 and ranged from 4,575 to 31,253. Across all articles, 61,117 articles were considered closed access whereas 85,298 had some form of OA. Specifically, 18,032 articles were classified as hybrid gold, 9,261 were classified as green, and 58,005 were classified as bronze.

In a simple generalized linear model analyses with access as the only independent variable, we found that hybrid gold articles had an average of 31.1 (30.6 - 31.5; 95% C.L.) citations, compared to 13.3 (13.2 - 13.4) citations for closed access articles. Bronze access articles averaged 35.9 (35.6 - 36.2) citations, while green access articles averaged 19.3 (18.9 - 19.7) citations. All categories of access were statistically different from one another (Wald *z* statistic with Bonferroni correction, all *z* > 16.411, all *p* < 0.0001).

Our full model indicated that, in addition to access type, all other variables included in the model had significant relationships with citation counts. Variation in the log number of citations due to the specific journal an article was published in had a standard deviation of 0.27. Similarly, variation in the log number of citations due to the biological sub-field in which an article was published had a standard deviation of 0.15. Moreover, we found that fixed-effects variables all interacted with access type to influence citation counts (Table 4). For example, the model suggested that hybrid gold access generated more citations than the other three access types when articles had few authors, but generated fewer citations than other access types when articles had many authors. Thus, to explore potential non-linearities in the relationship between number of authors and number of citations, as well as the interaction between number of authors and access type, we binned the number of authors into the following discrete categories: 1, 2, 3-4, 5-8, 9-16, 17-32, 33-64, 65-128, 129-256, and >257 authors. However, we note that only 118 (0.08%) articles had more than 64 authors. A likelihood-ratio test indicated that categorizing the author-count variable in this way significantly improved the model (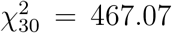, *p* < 0.0001). Therefore, we used this categorized variable in our full model analysis. The model indicated that with only a single author, hybrid gold generated 2.86 (3.66 - 2.24; 95% C.L.), 2.25 (2.72 - 1.86; 95% C.L.), and 2.08 (2.77 - 1.57; 95% C.L.) times as many citations as closed access, bronze, and green access types respectively (Fig. 2A; all *z* > 6.812, all Bonferroni-adjusted *p* < 0.0001). With only a single author, green and bronze also generated significantly more citations than closed access (*p* = 0.0121 and 0.0061, respectively), but differences were relatively small: 1.38 (1.05 - 1.81) and 1.27 (1.04 - 1.54) times as many citations, respectively. With one author, green and bronze access types were not significantly different from each other (*p* = 1.000). This ranked pattern in citations as a function of access was generally maintained between 2 and 32 authors, with hybrid gold generating the most citations, followed by green/bronze, and last by closed access (Fig. 2A). Differences among OA types were not always statistically significant, but OA types always generated significantly more citations than closed access over this author-count range (all *z* > 4.18, all *p* < 0.0002). Above 33 authors, differences among access types were more variable but typically not significantly different (Fig. 2A).

**Figure 2.**
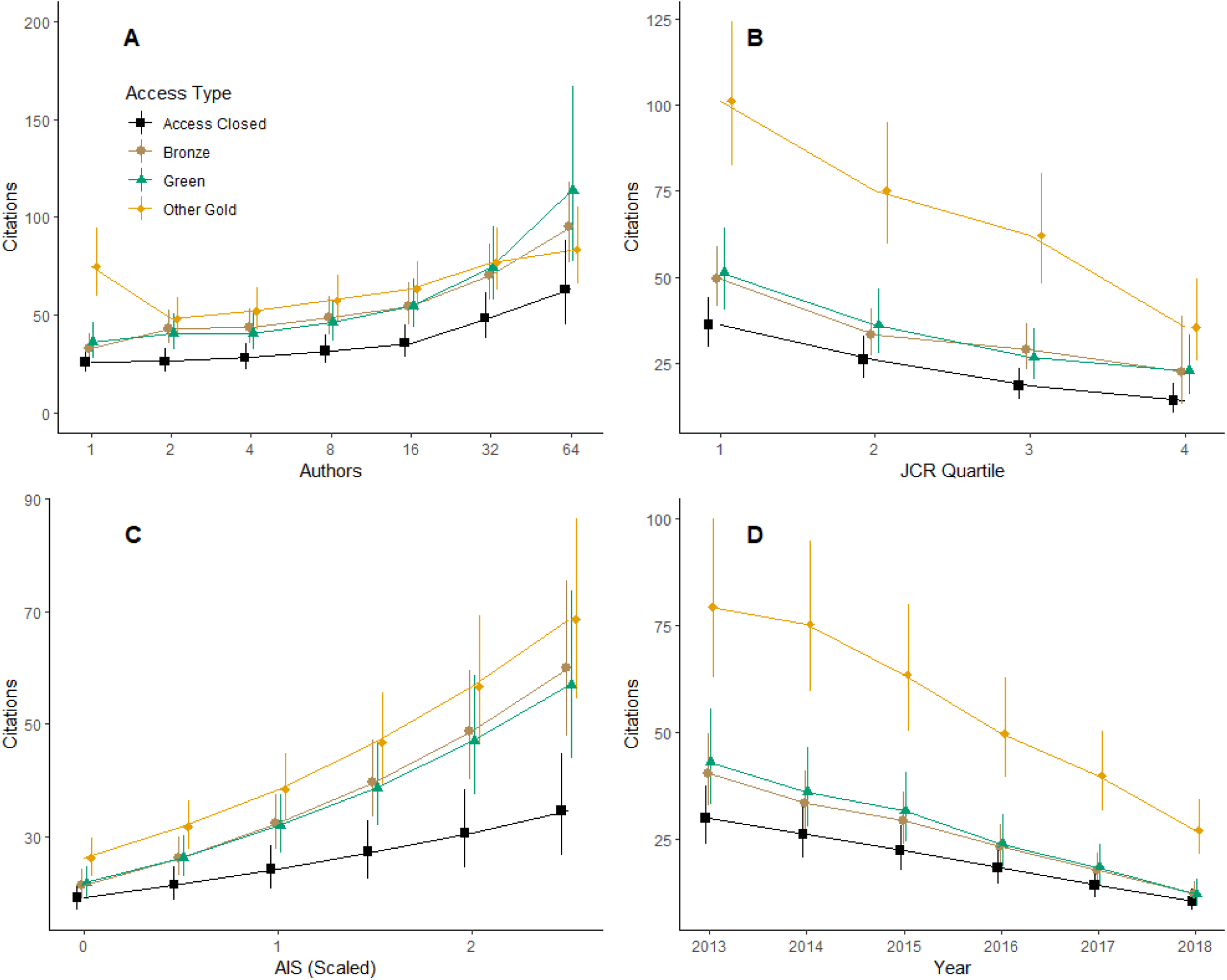
Citations as a function of various model parameters and access type. Access type is color coded to match the named color scheme (e.g. green is indicated in green), except closed access which is indicated in black. **A.** Interaction between access type and author count. The number of authors was treated as categorical. Only values under 64 are plotted to emphasize relationships in the majority of the dataset. **B.** Interaction between access type and JCR quartile. JCR Quartile of 1 represents the highest impact journals. **C.** Interaction between access type and scaled (standardized) AIS values. **D.** Interaction between access type and year.

**Table 4.**
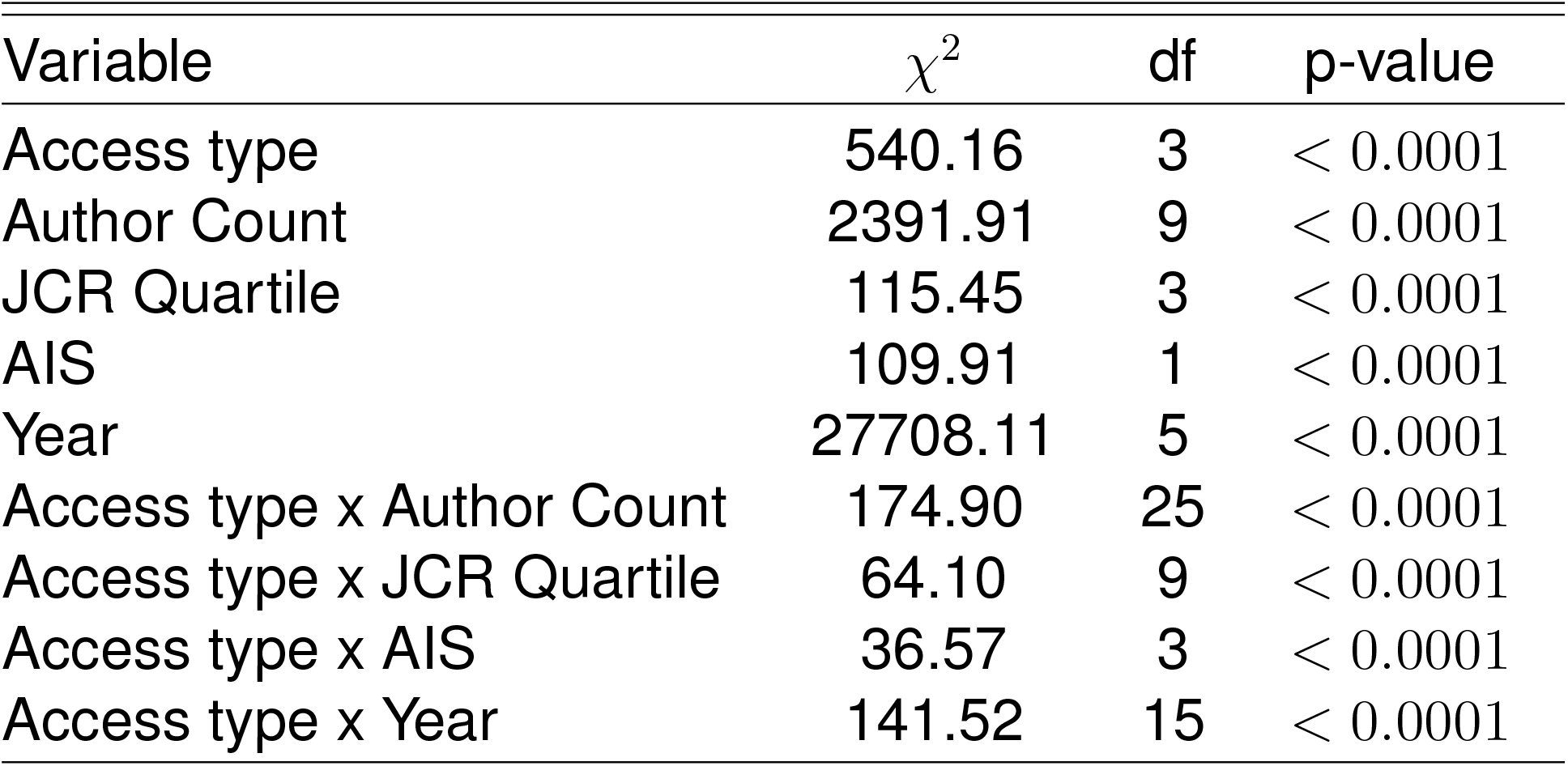
Results of analysis of full dataset. Type II Wald Chi-square tests were used. Thus, significance tests of interactions terms were marginal, but significance tests of main-effects terms were marginal excluding all interaction terms.

| Variable | $\chi^2$ | df | p-value |
| --- | --- | --- | --- |
| Access type | 540.16 | 3 | < 0.0001 |
| Author Count | 2391.91 | 9 | < 0.0001 |
| JCR Quartile | 115.45 | 3 | < 0.0001 |
| AIS | 109.91 | 1 | < 0.0001 |
| Year | 27708.11 | 5 | < 0.0001 |
| Access type x Author Count | 174.90 | 25 | < 0.0001 |
| Access type x JCR Quartile | 64.10 | 9 | < 0.0001 |
| Access type x AIS | 36.57 | 3 | < 0.0001 |
| Access type x Year | 141.52 | 15 | < 0.0001 |

Although the full model indicated significant interactions between JCR quartile and access type, as well year of publication and access type (Table 4), the general pattern of hybrid gold > green/bronze > closed access in terms of number of citations held across JCR Quartiles, scaled AIS, and year of publication (Fig. 2). However, the differences among the 4 types of access decreased with higher JCR Quartiles (Fig. 2B), lower scaled AIS scores (Fig. 2C), and year of publication (Fig. 2D). By the 4th quartile, differences between hybrid gold and bronze, and between bronze and closed access were no longer statistically significant (*z* = 1.80, 1.84; *p* = 0.43, 0.40, respectively). Similarly, at scaled AIS values < 1.5, differences between bronze/green and closed access were not significantly different (all *z* > 2.66, all *p* > 0.05). Finally, from 2016 to 2018, differences between green and closed access were not significantly different (all *z* > 2.56; all *p* > 0.062), and in 2018 differences between bronze and closed access were not significantly different (*z* = 2.02; *p* = 0.26).

Our matched analysis included 28,081 journal articles from 129 journals across the same 12 fields within biology (Table 2). The number of records per journal averaged 218 and ranged from 2 to 8,624. Records across research fields averaged 2,340 and ranged from 7 to 8,624 as some fields were only represented by a single journal. Across all articles, 23,598 were considered closed access whereas 4,483 were classified as hybrid gold.

Our model of the matched analysis dataset indicated that, in addition to access type, all other variables included in the model had significant relationships with citation counts, but moreover, the variables all interacted with access type to influence citation counts, except for year (Table 5). Unlike with the full dataset, treating author count as a categorical (i.e., binned) variable did not significantly improve the model (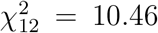, *p* = 0.56). The model indicated that with only a single author, hybrid gold generated 1.26 (1.136 - 1.4; 95% C.L.) times as many citations as closed access (Fig. 3A; *z* = 4.339, Bonferroni-adjusted *p* < 0.0001). The differences between hybrid gold and closed access decreased with increasing number of authors until 16 authors, at which point differences were not statistically significant (Fig. 3A; *z* = 1.823, Bonferroni-adjusted *p* = 0.068). Above ~60 authors, differences between access types were once again significant, with closed access generating more citations than hybrid gold (Fig. 3A; *z* = −2.290, Bonferroni-adjusted *p* = 0.0220).

**Figure 3.**
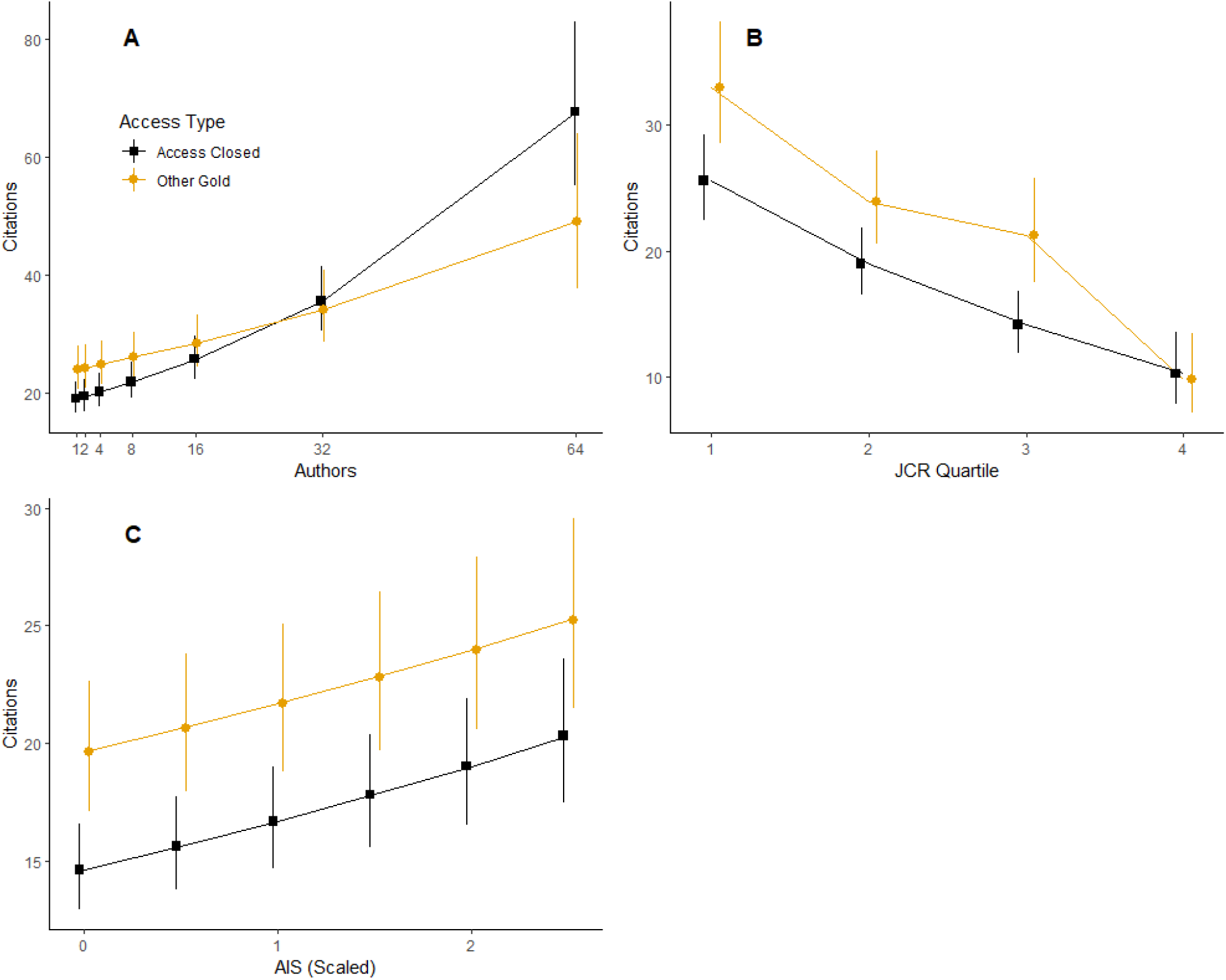
Citations as a function of various variables and access type from same issue of journal. Access type is color coded to match Figure 2. **A.** Interaction between access type and author count. **B.** Interaction between access type and JCR quartile. JCR Quartile of 1 represents the highest impact journals. **C.** Interaction between access type and scaled (standardized) AIS values. Note: Access type by Year is not shown here because this interaction term was not significant in this analysis (see Table 5).

**Table 5.**
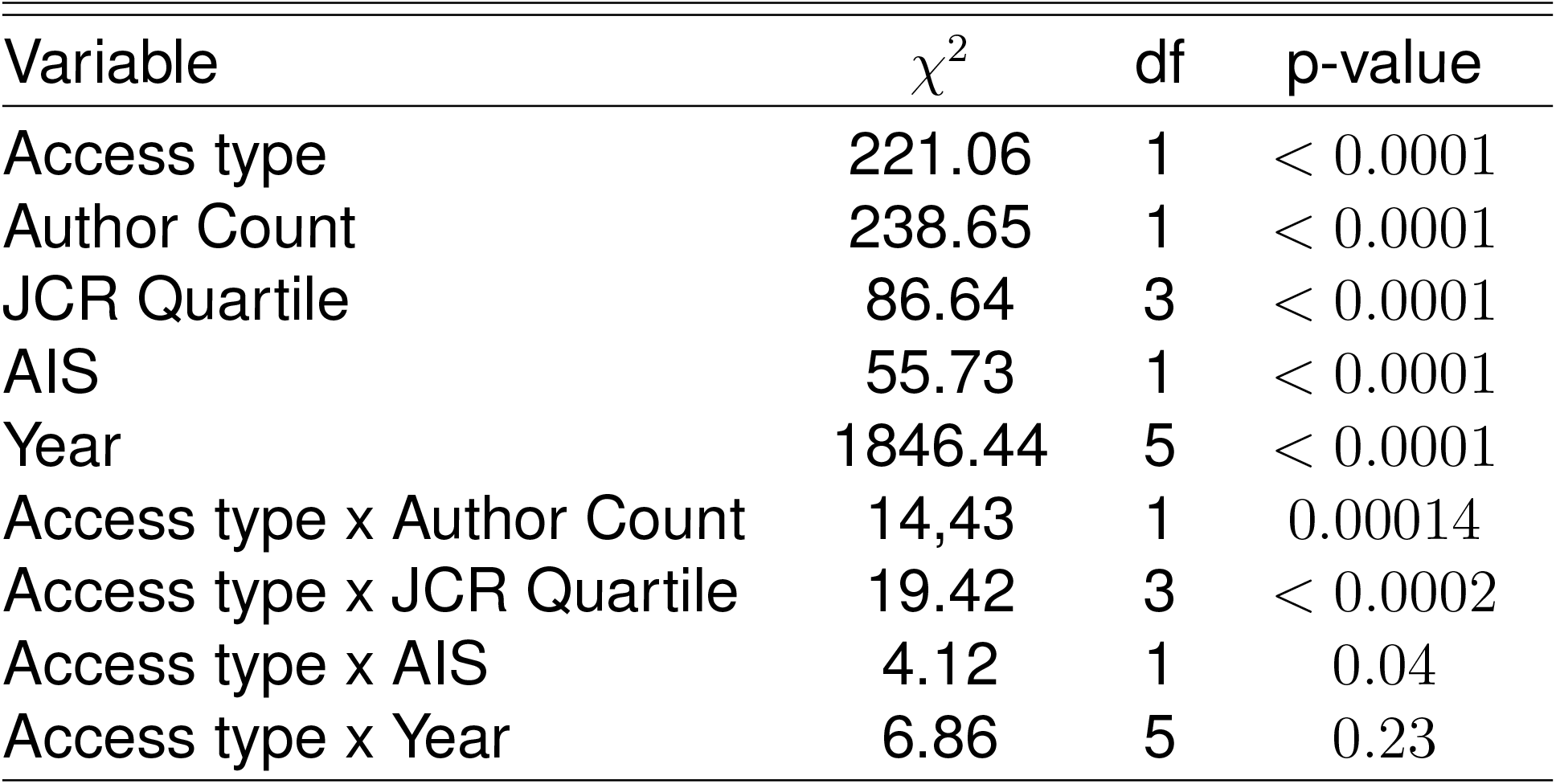
Results of analysis of matched dataset. Type II Wald Chi-square tests were used. Thus, significance tests of interactions terms were marginal, but significance tests of main-effects terms were marginal excluding all interaction terms.

| Variable | $\chi^2$ | df | p-value |
| --- | --- | --- | --- |
| Access type | 221.06 | 1 | < 0.0001 |
| Author Count | 238.65 | 1 | < 0.0001 |
| JCR Quartile | 86.64 | 3 | < 0.0001 |
| AIS | 55.73 | 1 | < 0.0001 |
| Year | 1846.44 | 5 | < 0.0001 |
| Access type x Author Count | 14.43 | 1 | 0.00014 |
| Access type x JCR Quartile | 19.42 | 3 | < 0.0002 |
| Access type x AIS | 4.12 | 1 | 0.04 |
| Access type x Year | 6.86 | 5 | 0.23 |

Similar to the analysis of the full dataset, and despite the interaction, the general pattern of hybrid gold > closed access, in terms of number of citations, held across JCR Quartiles and scaled AIS values (Figs. 3B,C). As with the full dataset, the differences between the two types of access decreased with higher-numbered JCR Quartiles (Fig. 3B) or lower scaled AIS scores (Fig. 3C). By the 4th JCR quartile, differences between hybrid gold and closed access were no longer statistically significant (Fig. 3B; *z* = −0.407; *p* = 0.68). Finally, we observed significantly greater number of citations for hybrid gold compared to closed access at all scaled AIS values, although differences decreased very slightly with increasing AIS values (Fig. 3C; all *z* > 3.677, all *p* > 0.0002).

Our APC analysis included 17,542 journal articles from 152 journals across 11 fields; the field of Biochemistry and Cellular Biology was comprised of articles from a single journal, and hence was removed from the analysis due to limited sample sizes that impacted model convergence. The number of records per journal averaged 116 and ranged from 1 to 2,649. Records across research fields averaged 1594 and ranged from 270 to 4,501. In a simple analysis of the relationship between number of citations and APCs, we found that for each standard deviation increase in APC (about $1500), we observed a 19.7% (17.7% - 21.9%; 95% C.L.) increase in the number of citations (*p* < 0.0001). However, we also found that for each $1000 dollar increase in APC, there was about a 1 unit (0.93 ± 0.03; ± 95% C.I.) increase in AIS (standard linear regression; *p* < 0.0001; *r*^2^ = 0.21; note that AIS in the data ranged from 0.17 to 20.8). After statistically controlling for AIS, JCR quartile, author count, and year, we found that the main effect of APC was not significant, but that there were significant interactions between APC and year, as well as APC and scaled AIS values (Table 6). Specifically, increasing APC resulted in slight increases in number of citations at low AIS, but almost no increase in number of citations at high AIS (Fig. 4A). Similarly, increasing APCs resulted in negligible to a slight increase in number of citations for all years except 2017, in which cases increasing APCs resulted in a decrease in the number of citations (Fig. 4B).

**Figure 4.**
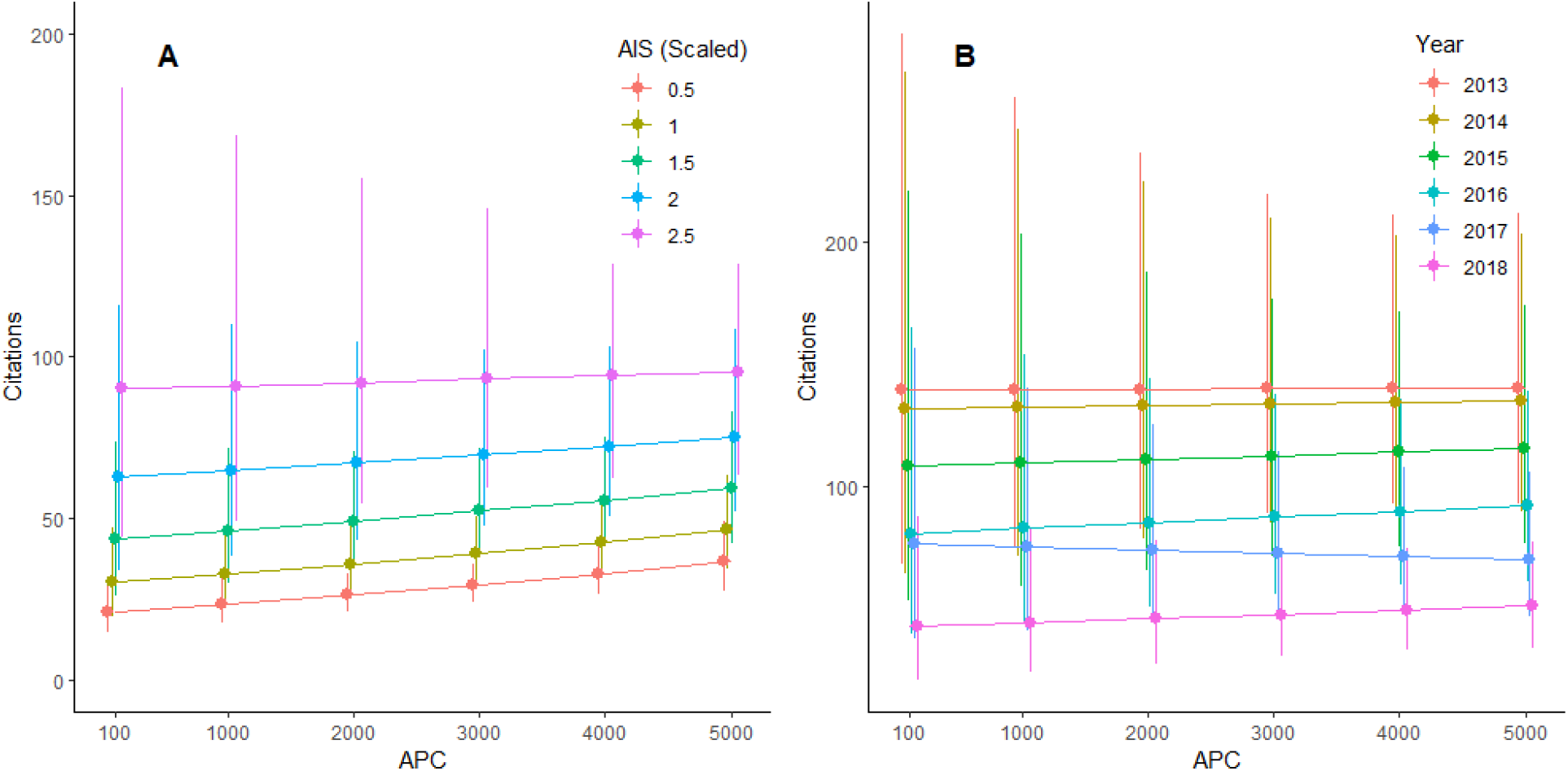
Citations in hybrid gold articles as a function of APC and AIS or year. **A.** Interaction between APC and scaled (standardized) AIS values. **B.** Interaction between APC and Year of Publication.

**Table 6.**
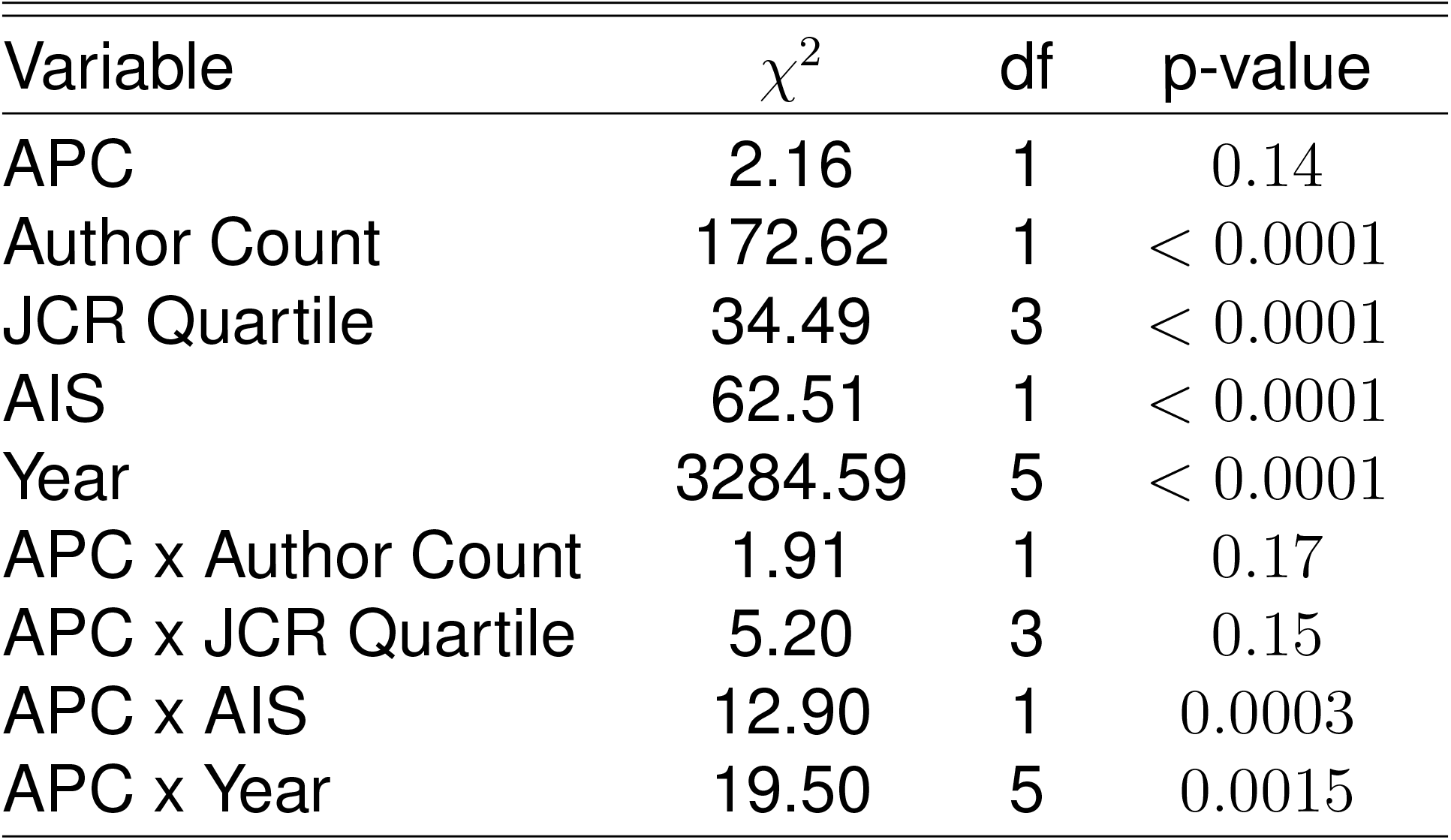
Results of analysis of APC dataset. Type II Wald Chi-square tests were used. Thus, significance tests of interactions terms were marginal, but significance tests of main-effects terms were marginal excluding all interaction terms.

## Discussion and Conclusion

All OA types—gold, green, and bronze—yielded a significant citation advantage relative to closed access articles published in hybrid journals (Figure 2). We did find a more pronounced advantage for hybrid gold articles than bronze or green articles, suggesting that paying an APC to make an article freely available provides a greater comparative citation advantage (but see further discussion of the APC charges below). The exceptions to this pattern appear to be articles with a large number of authors (>~32 authors), and those published in relatively low impact journals (i.e., those with AIS scores close to 0 and/or in the lowest JCR quartile). We observed the same general patterns when restricting our analyses to compare only hybrid gold and closed access articles that were published in the same issue of a journal (Figure 3).

When publishing under hybrid gold access models, journals with higher AIS scores (i.e., higher impact) tend to have higher APCs [Budzinski et al., 2020]. Thus, paying the higher APCs associated with higher impact journals may result in more citations, a pattern we recovered support for in our APC analysis. However, after controlling for the effect of journal impact quantified by AIS, higher APCs had minimal effects on citation counts. Disentangling the effects of journal impact when trying to assess whether paying an APC adds additional citations has proven difficult, as authors may prioritize making their more impactful work open [Craig et al., 2007]. Our results were consistent with Piwowar et al. [2018], who found an OA citation advantage primarily driven by hybrid gold publishing.

Our results indicate that paying an APC for gold OA in a hybrid journal or self-depositing at no cost to the authors is a tradeoff between time and money. Opting for the gold route by paying an APC allows an article to be freely available immediately upon publication, increasing the potential audience size by removing barriers to reader access while the research is new, likely increasing the attention the article receives. Indeed, our results suggest that publishing gold OA article in a hybrid journal maximizes citations in most scenarios. Additionally, choosing to publish gold OA avoids the embargo period imposed by publishers that authors face when choosing to publish green OA, which may last six months to a year post-publication. That said, our results do indicate that self-archiving also confers an (albeit less pronounced) citation advantage; therefore, if funds are not available it is still advantageous for authors to deposit their works in repositories (Figure 5).

**Figure 5.**
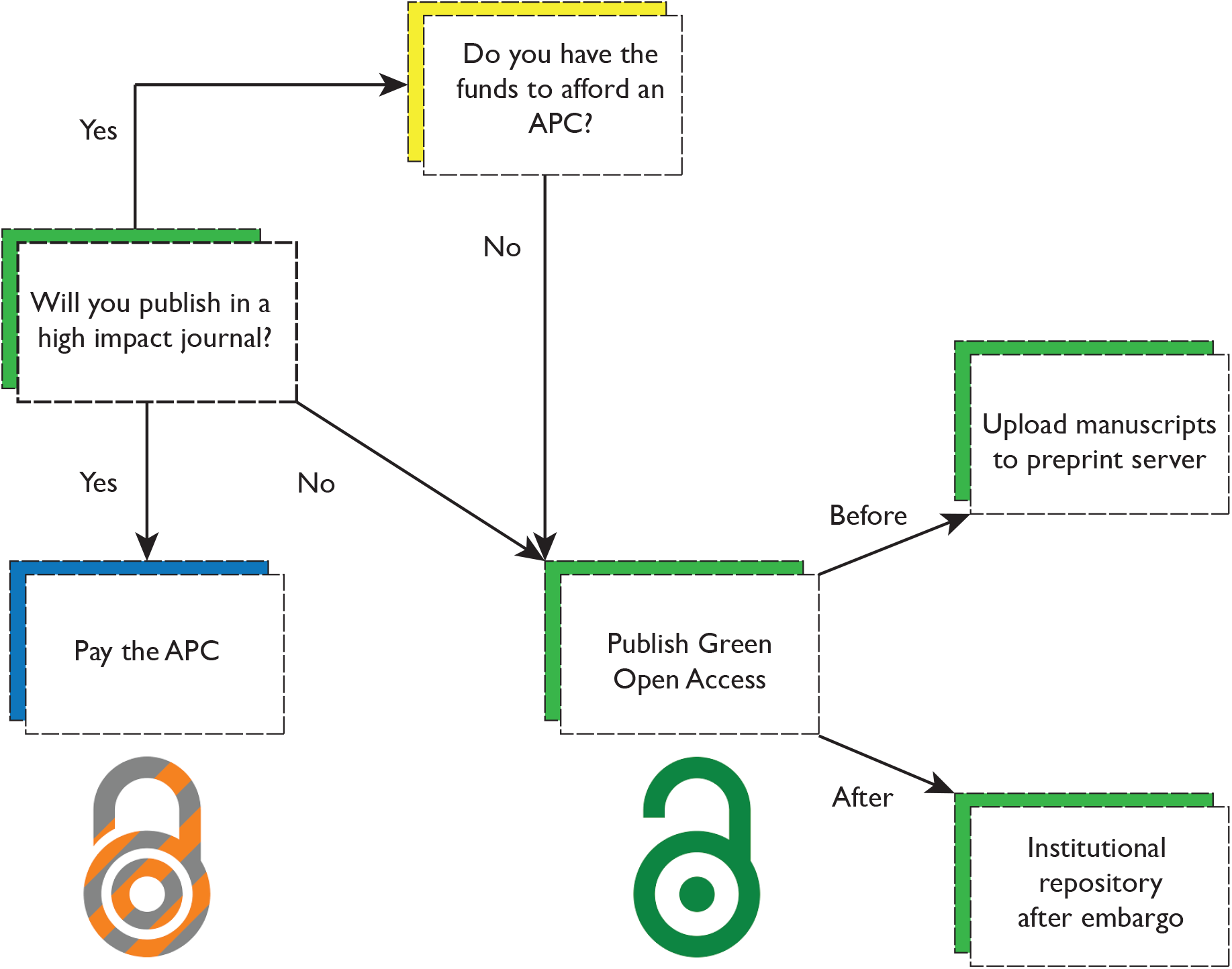
What to consider when publishing open access. Authors have many junctures between article submission and appearance in a journal when they can make their research available while also saving money. After deciding which journal is a good fit for their reserach, if the journal is low impact, authors can opt for the green route without sacrificing many potential citations (although a low impact journal likely will not have a prohibitively expensive APC). If the journal is of high impact, whether author(s) should pay an APC comes down to their funding—if they have limited funds, they should forgo paying an APC and opt for the green route as that will be likely to garner more citations than publishing closed access. If authors go with the green route, they can submit their articles to a pre-print server before publication and archive in an institutional repository post-publication; the latter suggestion also applies to articles published closed access older than two years.

Publishers are aware of this tradeoff, as evidenced by the mainstreaming of the hybrid model, the rise of APCs themselves [Budzinski et al., 2020], and increased restrictions on self-archiving [Gadd and Troll Covey, 2019]. Though many authors have noticed higher APCs within the same journals over time [Khoo, 2019], historical data on APCs is difficult to find. In 2019, then-current APCs were published for a selection of journals listed in the Directory of Open Access Journals [Krzton, 2019]. In a 4-year time frame (between 2019 and 2022), the APCs for all but one journal became more expensive, and one title (PLoS Biology) increased its APC by nearly 77% (Table 1). While these figures are from gold (fully OA) journals rather than hybrid journals, they do reflect the trend of increasing APCs, and several of those journals are controlled by commercial publishers that also have substantial hybrid journal offerings within biology. Along with rising APCs, publishers have increased restrictions on the conditions of self-archiving one’s work, particularly by adding embargoes on green OA deposits [Gadd and Troll Covey, 2019]. Beyond putting authors in the difficult position of having to choose between allocating funds to new research or publishing their existing work gold OA, these new barriers threaten to drive further inequality between the Global North and South, which has spurred a growing movement to eliminate APCs altogether [Alperin, 2022, Peterson et al., 2019, Alizon, 2018, Mekonnen et al., 2022].

In light of the foregoing discussion, we offer the following recommendations to authors submitting a manuscript to a biology journal (visualized in Fig. 5):

### • Choice of journal should not be dependent on open-access status

Due to the widespread adoption of OA through a variety of channels, most authors today have the option of publishing OA, regardless of target journal. Authors should choose the best-fit journal for their research according to their preferred criteria, separate from the issue of when and how to make their work open. The only exception would be if a research sponsor mandates gold OA, in which case authors could not submit to subscription journals that did not at least offer a hybrid option.

### • If a research sponsor requires gold OA, their funding should cover the APC

Authors should review the terms of sponsored research agreements closely to see whether any resulting publications are required to be OA. Some sponsors specify immediate public access to the version of record via gold OA, in which case authors should request that sponsor cover the APC if publication fees are not already written into the grant.

### • Authors should save the final accepted manuscript version for later deposit in institutional repositories

Many journals that permit green OA via deposit into an institutional repository still prohibit deposit of the publisher’s PDF with all journal formatting and type-setting applied. When the final version of the manuscript has been approved by the journal editors and all authors, at least one author should retain that version in manuscript form to deposit into an open repository. If the journal requires an embargo period, contact repositories to see whether an immediate deposit is possible with an embargo that will automatically expire on a certain date. This eliminates the need for authors to personally keep track of when they can self-deposit and also minimizes the chances of misplacing the manuscript file in the meantime.

### • Consider depositing all closed access articles over two years old

Green OA articles were found to have a citation advantage in this study and others [Ottaviani, 2016]. The more restrictive commercial publishers typically set their embargo period for self-deposit in a repository at two years, with most others allowing for self-deposit after one year or even six months. Any article published two years ago or more is almost certainly eligible to be deposited into an open repository. Authors can leverage the OA citation advantage for these older articles at no cost to themselves, and some institutions may provide assistance with deposit through their scholarly communication units and/or libraries.

### • Ensure when paying an APC that the article will receive a recognized open license

When authors pay an APC, it is important to verify that the article will be published under a “CC-BY” or similar open license and that the license will be clearly listed either on the journal page or in the text of the article itself. This guarantees that authors who pay APCs are providing gold, rather than bronze, OA to their work.

### • Check whether your institution has a nonexclusive right to deposit prior to publication

Some universities have adopted policies that assert a nonexclusive right to distribute scholarly work by affiliated personnel on behalf of the authors. These policies are designed to supersede publisher embargoes on self-deposit and may allow authors to open their articles via the green route immediately upon acceptance, without paying an APC.

## Data Availability

Raw data and scripts used to replicate the analyses conducted here were archived in a GitHub repository and are available on Zenodo: https://doi.org/10.5281/zenodo.7416222. Additionally, the models themselves have been saved as R objects and deposited in the Auburn University institutional archive AUrora: https://aurora.auburn.edu/handle/11200/50478.

## Acknowledgments

This project began as a class assignment for the Fall 2020 iteration of the BIOL 6800 course at Auburn University. LSS served as instructor of the course and ADC served as the graduate teaching assistant. We would like to thank those students for their contributions to this work many of whom appear as co-authors here.

